# TFL1 homolog integrates temperature signals to control timing of developmental transitions in woodland strawberry

**DOI:** 10.64898/2026.10.20.760361

**Authors:** Sergei Lembinen, Feng Wang, Quan Zhou, Teng Zhang, Guangxun Fan, Paula Elomaa, Timo Hytönen

## Abstract

- In overwintering plants of temperate climates, inflorescence and flower buds commonly form in autumn under cool temperatures, while flowering occurs the following spring. However, how cool temperatures regulate inflorescence and flower development remains poorly understood.
- We used woodland strawberry (*Fragaria vesca* L.), a perennial Rosaceae herb, to investigate the effects of cool temperature on reproductive development. We characterized temperature-dependent developmental patterns at macroscopic and microscopic scale and combined time- and stage-specific gene expression analyses.
- We show that cool temperature accelerates floral transition but subsequently slows inflorescence and flower development, causing increased inflorescence branching, higher floral organ numbers, and berry fasciation. These effects were restricted to the apical inflorescence and flower meristems, whereas higher-order meristems remained unaffected. Expression analyses revealed stage- and temperature-dependent changes in the abundance and spatial distribution of *TERMINAL FLOWER1* (*FvTFL1*) encoding the floral repressor. Loss of functional *FvTFL1* abolished temperature-dependent differences in floral transition, inflorescence branching, and floral organ numbers.
- Our results establish *FvTFL1* as a major regulator of temperature-responsive reproductive development. The stage-specific activity of *FvTFL1* provides a mechanism through which woodland strawberry integrates temperature signals to balance developmental plasticity of the primary growth axis with the robustness of lateral branches.

## Introduction

In plants growing in temperate regions, floral transition and anthesis are often separated between the seasons. In plants with such life history, inflorescence and flower buds are established during autumn and early winter, while flowers emerge during the following spring (Schnablová et al. 2021; Penfield 2024). Cool temperature acts as a primary cue regulating the transition from vegetative to reproductive development. Prolonged exposure to low temperatures, known as vernalization, promotes floral transition. Although vernalization is generally regarded as a response to winter conditions, in many species the upper temperature limit for effective vernalization can be relatively high (Lang 1965; Wollenberg and Amasino 2012; Duncan et al. 2015; O’Neill et al. 2019; Fan et al. 2022). As a result, many species fulfill their vernalization requirement and initiate flower buds before the onset of winter.

The requirement for vernalization is mediated by distinct floral repressors across different species (Bouche et al. 2017). In winter annual and perennial Brassicaceae, MADS-box genes orthologous to *FLOWERING LOCUS C* (*FLC*) in Arabidopsis confer vernalization requirement and seasonal flowering habit (Wang et al. 2009, Madrid et al. 2021, Zhai et al. 2024). Instead, in strawberries and sugar beet cool temperature requirement for flowering is conferred by the members of Phosphatidylethanolamine-Binding Protein (PEBP) family, homologs of *TERMINAL FLOWER1* (*TFL1*) and *FLOWERING LOCUS T* (*FT*), respectively (Pin et al. 2010, Koskela et al. 2012, 2016, Fan et al. 2022). Under low temperatures, *FLC* is progressively repressed through chromatin modifications and antisense transcription (Pandey et al. 2021, Sung et al. 2006; Swiezewski et al. 2009). Variation in the initial expression level and the downregulation dynamics of *FLC* orthologs contributes to interspecific and intraspecific differences in temperature requirements and duration of vernalization necessary for floral transition (Albani et al. 2012; Kemi et al. 2013; Duncan et al. 2015; O’Neill et al. 2019; Hepworth et al. 2020, Calderwood et al. 2021, Wunder et al. 2023, Zhu et al. 2023). Similarly in woodland strawberry, gradual downregulation of *FvTFL1* enables floral transition in autumn (Koskela et al. 2012). Cool temperatures below ∼13°C downregulate *FvTFL1* independently of photoperiod, whereas under warmer temperatures, short days are required for its downregulation (Rantanen et al. 2015). In natural accessions, higher *FvTFL1* expression levels delay floral transition and flowering (Zhou et al. 2026).

Although the role of vernalization in floral transition has been extensively studied, surprisingly little is known about the regulation of inflorescence and flower bud development under vernalizing temperatures. Inflorescence formation involves developmental transition of vegetative meristems into inflorescence meristems that ultimately produce flower meristems, the process also called meristem maturation (Park et al. 2014). The timing and rate of these transitions are particularly important in agricultural crops because they may underlie variation in inflorescence architecture and yield (Park et al. 2014; Eshed & Lippman 2019), yet the regulatory mechanisms controlling inflorescence and flower development under cool temperatures remain poorly understood. Addressing this gap becomes particularly important in the context of climate change because temperature variation between seasons may underlie changes in yield of winter annual and perennial crops (Døving and Måge 2001, O’Neill et al. 2019, Lu et al. 2022, Wen et al. 2026). In summer annual Arabidopsis, leaf-produced FT integrates photoperiod and temperature signals to regulate inflorescence development through interactions with *LFY* and *TFL1* (Lee et al. 2019; Huang et al. 2026). TFL1 maintains inflorescence meristem identity and indeterminacy by repressing the expression of flower-promoting transcription factors such as *LEAFY* (*LFY*) and *APETALA1* (*AP1*), whereas *tfl1* mutant generates a small inflorescence with a terminal flower (Alvarez et al. 1992; Zhu et al. 2020). Similarly, *fvtfl1* mutant of woodland strawberry that flowers perpetually in long days exhibits reduced inflorescence branching that is photoperiodically regulated via *FvFT1* (Lembinen et al. 2023). In contrast, in winter annual and perennial Brassicaceae, and seasonal flowering woodland strawberry, inflorescence formation in autumn occurs largely in the absence of *FT* expression in leaves (Koskela et al. 2012; González-Souárez et al. 2025). Nevertheless, *FT* orthologs are required for normal inflorescence and flower development in perennial *A. alpina*, which suggests that they may function in tissue specific rather that systemic manner (Sashidhar and Coupland 2026). Furthermore, incomplete downregulation of floral repressors such as FLC during vernalization leads to reduced inflorescence branching and floral reversion phenotypes (Lazaro et al. 2018; Lu et al. 2022), indicating that they continue to shape inflorescence and flower development even after floral transition.

Here we examined the effects of temperature on the timing of floral transition, inflorescence branching and flower development in woodland strawberry. We find that cooler temperatures accelerated floral transition but delayed inflorescence and flower development, resulting in extensive inflorescence branching and flower fasciation. We demonstrate that *FvTFL1* is largely responsible for this developmental heterochrony that is associated with temporal and spatial differences in its expression pattern under different temperatures.

## Results

### Cool temperature accelerates floral transition in woodland strawberry

To investigate the role of temperature in reproductive development of woodland strawberry, we subjected a seasonal flowering accession FIN56 to 17°C and 11°C (hereafter referred as warm and cool temperatures) in short days (12h/12h day/night) for six weeks. After each week, a subset of plants was moved to warm long days (20°C; 18h/6h day/night) where flowering was observed. The exposure to cool temperature strongly promoted floral transition as 44% of plants flowered after three weeks of treatment and a 4-week treatment induced flowering in all plants. However, after four weeks at warm temperature flowering was induced in only 16% of plants, and even after five weeks 12% of plants remained vegetative (**Figure 1A**). These observations were concordant with the number of leaves the plants initiated before switching to produce terminal inflorescence. At cool temperature, plants produced 7-9 leaves on the main axis before a terminal inflorescence, while after the warm treatment, they produced 9-13 leaves (**Figure 1B**). The differences in flowering time, i.e. days to the first open flower, however showed a different pattern. When moved to long days, plants exposed to treatments for four and five weeks flowered at the similar time (**Figure 1C**, *p = 0.79* week 4, *p = 0.78* week 5), whereas six weeks of cool temperature treatment delayed anthesis compared to the warm treatment (*p = 0.00013*). Therefore, although floral transition was advanced by cool temperature by at least a week, flower anthesis was not promoted or was even delayed by longer exposures to cool temperature compared to warm temperature treatments. In general, however, longer exposure to both warm and cool temperatures accelerated subsequent anthesis, indicating that the later flowering observed after six weeks of cool temperature was not caused by cessation of growth but rather a different developmental process.

**Figure 1.**
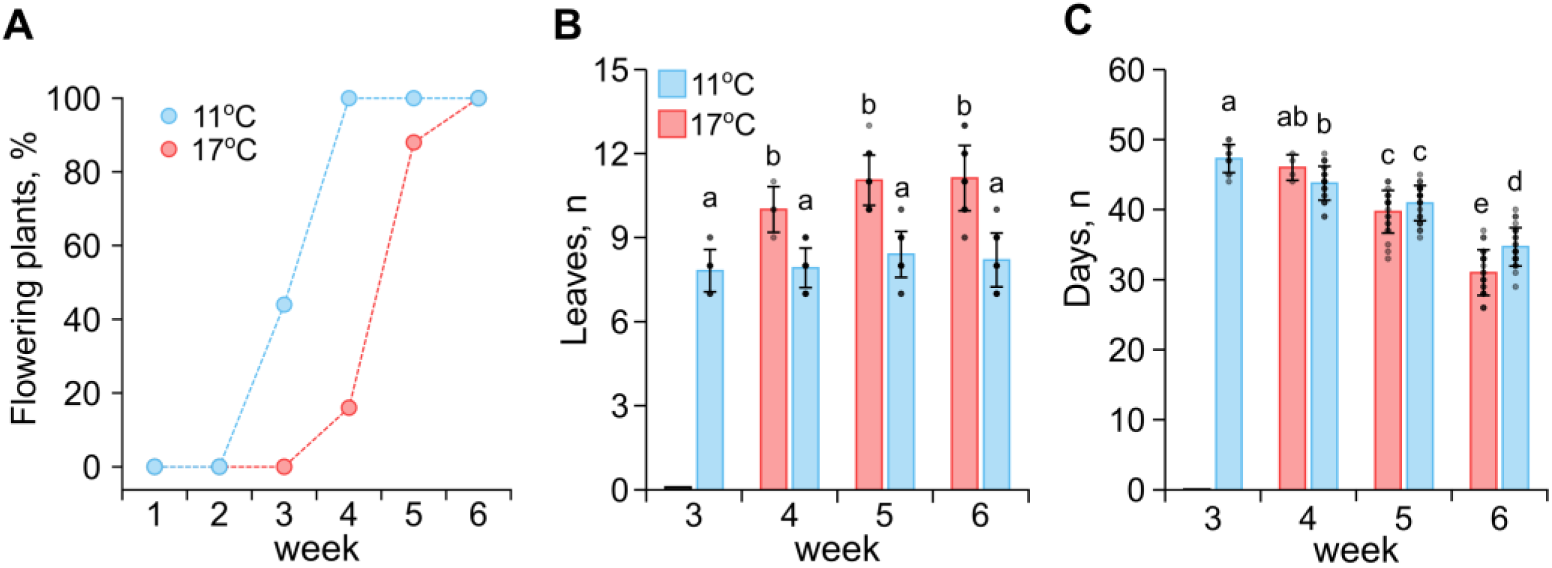
The effects of warmer and cooler temperatures on flowering time and plant architecture in woodland strawberry. **A.** Percentage of flowering plants after exposure to indicated temperatures for 1 – 6 weeks. **B.** Number of leaves before the terminal inflorescence after indicated treatments of 3 – 6 weeks. **C.** Flowering time as number of days to anthesis after the end of temperature treatments of 3 – 6 weeks. Different letters indicate statistically significant differences as determined by two-way ANOVA followed by Tukey’s HSD multiple comparisons test (*P* < 0.05; n = 11 - 25).

### Temperature regulates inflorescence branching in woodland strawberry

To further understand how temperature regulates reproductive development in woodland strawberry, we analyzed inflorescence architectures. The inflorescence of woodland strawberry consists of lateral dichasial cymose branches along the determinate primary axis (Lembinen et al. 2023). We found that the plants which flowered after the exposure to warm temperature for five and six weeks produced inflorescences consisting of 2 – 3 lateral dichasial branches and 15 – 25 flowers (**Figure 2A – C**). The exposure to extended periods of cool temperature, however, caused a gradual increase in inflorescence branches and flowers. Plants exposed to cool temperature for three weeks produced 2 – 4 lateral branches, whereas longer treatments allowed formation of inflorescences with up to six lateral branches and ultimately more flowers (**Figure 2A, B, D, E**). The effect of cool temperature on inflorescence branching was confined to the primary axis, and the lateral axes maintained their dichasial branching pattern, suggesting that their development is regulated differently. The increased number of lateral branches on the primary axis was previously shown to be associated with delayed termination of the apical inflorescence meristem (Lembinen et al. 2023). Therefore, even though the floral transition was promoted at cooler temperatures, the development of inflorescences was delayed resulting in increased branching.

**Figure 2.**
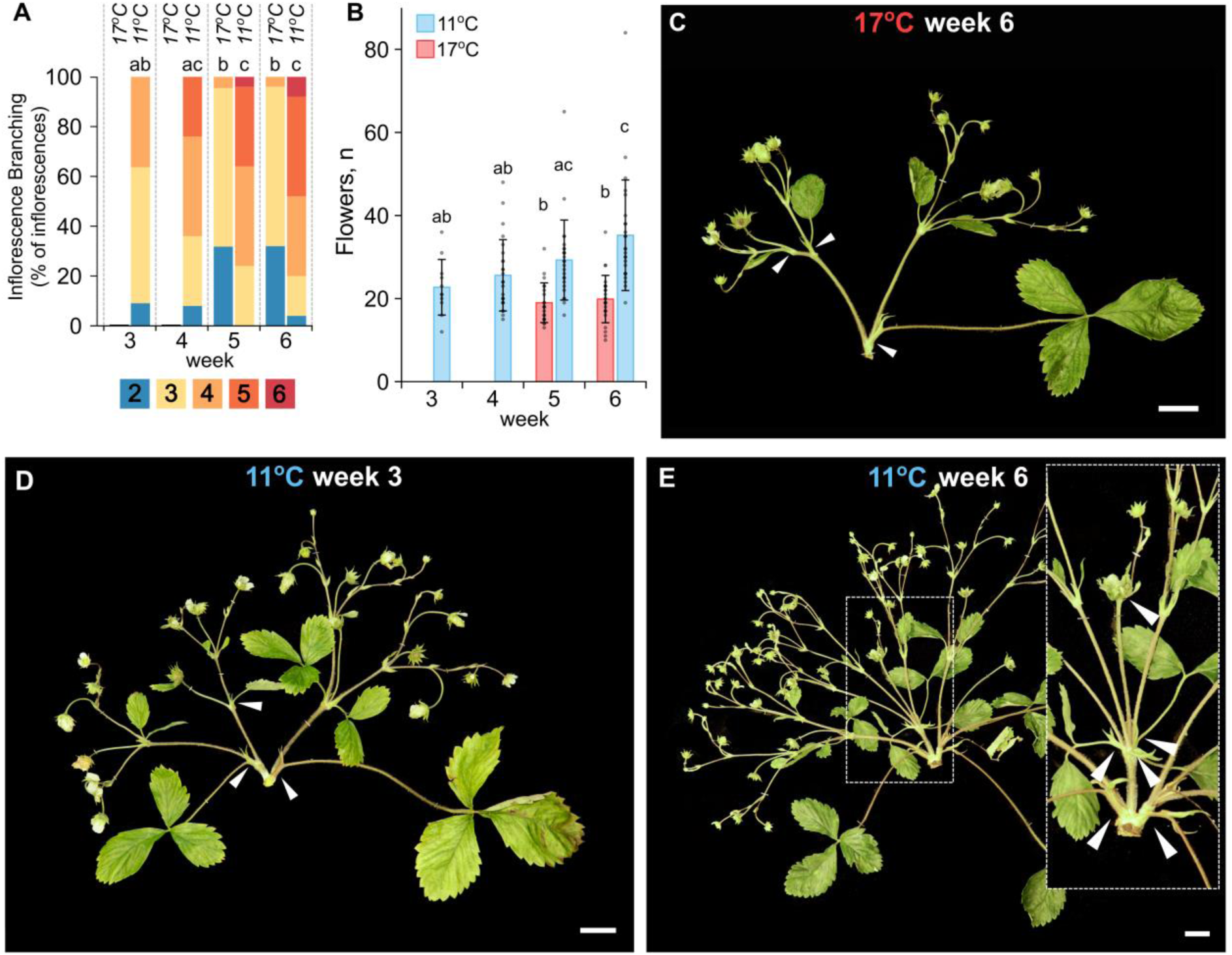
Temperature affects inflorescence branching and architecture in woodland strawberry. **A.** Percentage of inflorescences with indicated number of lateral branches after warm (17 °C) and cool (11 °C) temperature treatments of different durations. **B.** Number of flowers per inflorescence after warm and cool temperature treatments of different durations. Different letters indicate statistically significant differences as determined by two-way ANOVA followed by Tukey’s HSD multiple comparisons test (*P* < 0.05; n = 11 - 25). **C – E.** Examples of inflorescences produced after six weeks of warm temperature treatment (**C**), and three (**D**) and six weeks (**E**) of cool temperature treatment. White triangles indicate lateral branches on the primary growth axis. Scale bars are equal to 2 cm.

### Longer exposure to cool temperature induces flower and berry fasciation

Flowers of woodland strawberry typically consist of 5 episepals, 5 sepals, 5 petals, 20 – 25 stamens and about a hundred of carpels (**Figure 3A**). The primary flowers which originate from the terminating apical meristem are the largest and often show deviations in organ numbers (Darrow 1929, 1966). We found that the plants induced to flower under warm temperature for five and six weeks produced 5 – 7 sepals, 6 – 8 petals, 20 – 35 stamens and 120 – 170 carpels on their primary flowers (**Figure 3A – E**). The flower induction at cooler temperature significantly increased the number of organs in each flower whorl (**Figure 3B – J**). In addition, chimeric floral organs such as bract-like sepals, sepaloid petals or petaloid stamens were occasionally observed (**Supplemental Figure S1A – D**). Similarly to inflorescence branching the effect of cool temperature on flower development accumulated with the duration of the treatment, however, temporal patterns were specific to each floral whorl. For example, the number of sepals was not strongly increased by the treatments longer than four weeks (**Figure 3B**), whereas the exposure to cool temperature for up to five weeks increased the number of petals (**Figure 3C**). The number of stamens and carpels increased towards the longest six-week treatment (**Figure 3D and E**). In addition to increased number of all floral organs, the longer exposure to cool temperature altered the shape of the receptacle causing fasciation (**Figure 3K and L**). The fasciation of the receptacles significantly increased the width of the berries and as a result their fresh weight (**Figure 3M and N**). Similarly to the inflorescence development, the temperature effect was confined to the primary flower formed by the apical meristem. Flowers in the higher order branches formed by the newly established axillary meristems developed similar number of organs under both treatments (**Supplemental Figure S2**). These results highlight the differences in the regulation of developmental transitions in the apical and axillary reproductive meristems.

**Figure 3.**
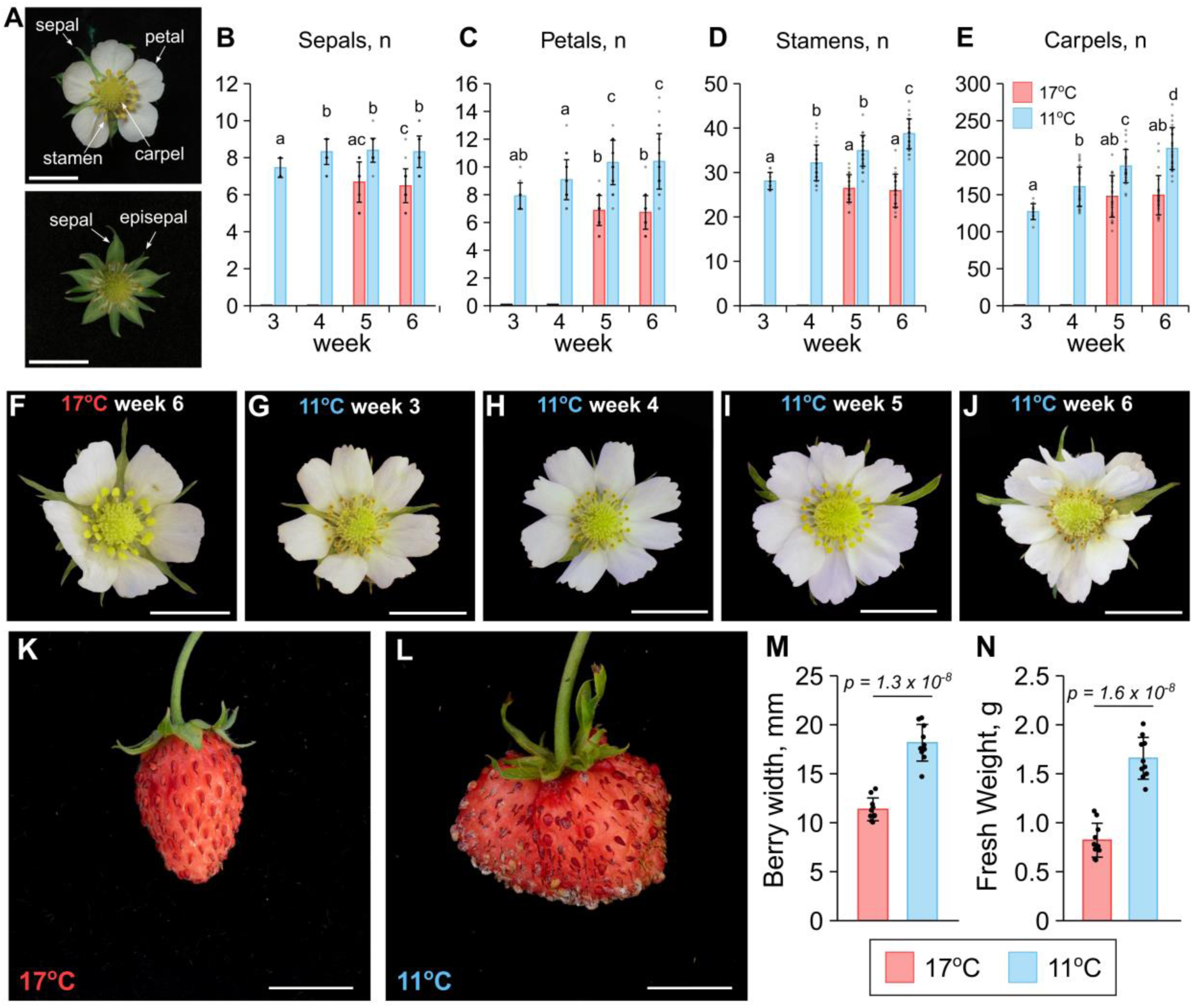
The effect of different temperature treatments on primary flowers and berries of woodland strawberry. **A.** Anatomy of woodland strawberry flowers. **B – E.** The number of sepals (**B**), petals (**C**), stamens (**D**), and carpels (**E**) developed on the primary flowers under warm (17°C) and cool (11°C) temperature treatments. Different letters indicate statistically significant differences as determined by two-way ANOVA followed by Tukey’s HSD multiple comparisons test (*P* < 0.05; n = 11 - 25). **F – J.** Examples of flowers developed after the temperature treatments of different duration. **K and J.** Berries developed from primary flowers after exposure to 17°C and 11°C for six weeks. Scale bars are equal to 1 cm. **M and N.** Width (**M**) and fresh weight (**N**) of the berries developed from primary flowers. Data are the mean ± SD (n = 10).

### Temperature fine-tunes the rate of meristem maturation in woodland strawberry

To understand how exposure to cool temperatures accelerated floral transition and promoted inflorescence branching and berry fasciation we exposed plants to warm and cool temperature treatments and collected meristems at weekly intervals (**Figure 4A and B**). Under warm temperature the apical meristems showed no significant morphological change over the first three weeks of the treatment. The visible expansion appeared after four weeks, and after five weeks the apical meristem was fully transformed into a floral meristem with sepal and petal primordia. After six weeks the flower bud was enclosed by sepals. When plants were grown under cool temperature, the expansion of the apical meristem was observed already after three weeks, but further development was delayed. Sepal primordia were observed at week 5, and even after six weeks of cool temperature treatment, only petal primordia emerged, corresponding to the developmental stage of the meristem after five weeks of warmer treatment. Interestingly, the first 2 – 3 sepals of woodland strawberry primary flowers were larger and more serrated than those developed later (**Supplemental Figure S3A - F**), suggesting a mixed identity between bracts and true sepals. The developmental differences were more evident in flowers developed under cool temperature with the first 2 – 3 sepal primordia developing farther from the center of the meristem than other sepal primordia (**Figure 4A and B**). This pattern of sepal formation likely contributed to the various degrees of receptacle fasciation observed under longer cool temperature treatments (**Supplemental Figure S3A-D**). Overall, we propose that the observed phenotypic differences in timing of floral transition, inflorescence branching, and flower ontogeny under different temperatures are due to the heterochrony in the apical meristem maturation. Our data indicates that cool temperature first promotes the floral transition (SAM to IM) but then delays subsequent IM to FM transition, and flower meristem patterning and termination.

**Figure 4.**
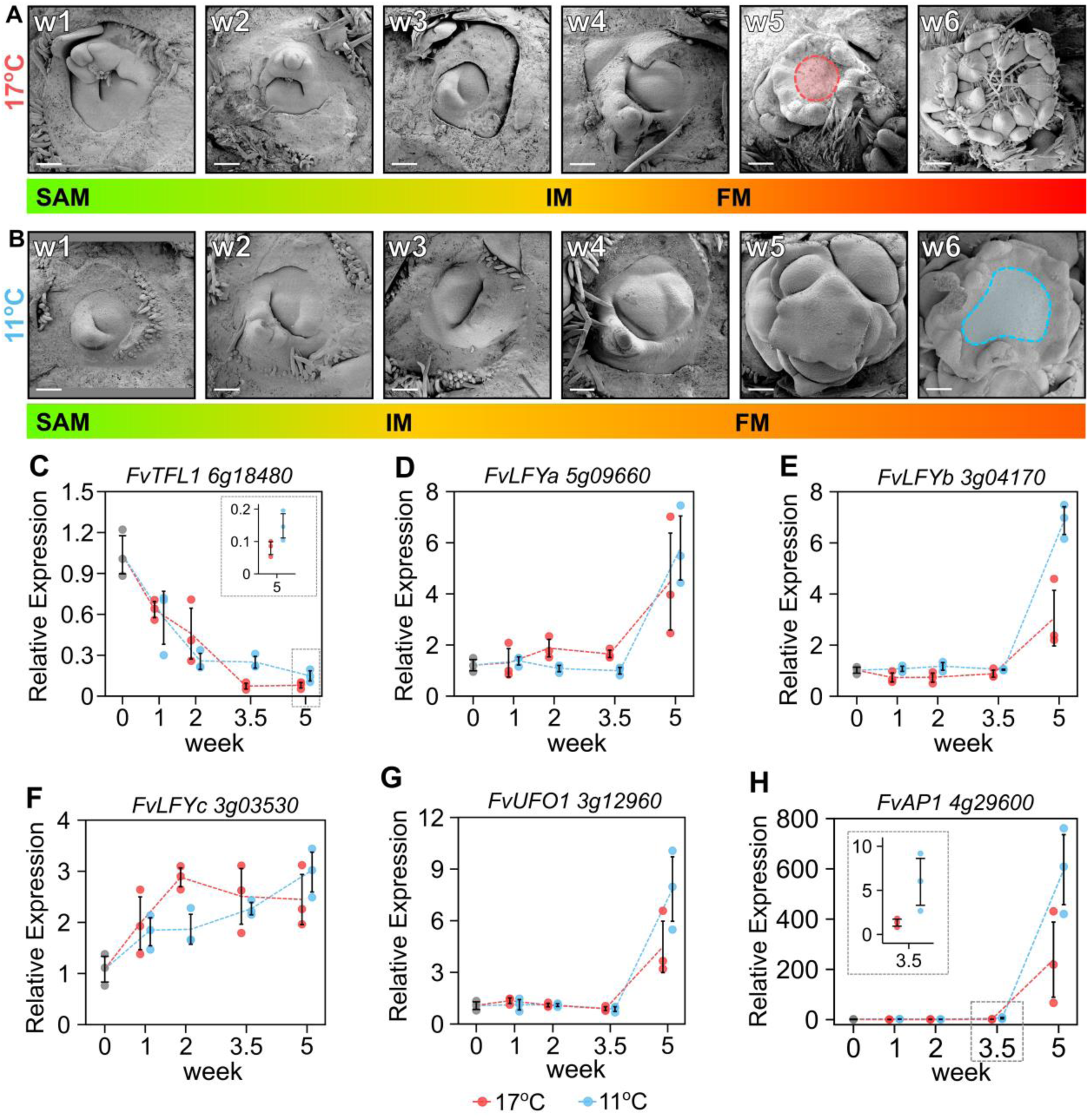
Heterochrony caused by the temperature treatments. **A - B.** SEM images showing weekly morphological changes in the meristems at warm (17°C) (**A**) and cool (11°C) (**B**) temperatures. Highlighted areas indicate the shape of the receptacle area that develops into a berry. Scale bars are equal to 100 µm. Color gradient bars under images represent meristem identity transitions. **C – H.** Temporal expression patterns of *FvTFL1* (**C**), *FvLFYa* (**D**), *FvLFYb* (**E**), *FvLFYc* (**F**), *FvAP1* (**G**), and *FvUFO1* (**H**) at warm (17°C, red) and cool (11°C, blue) temperatures. Error bars represent SD of three biological replicates (colored dots).

We then analyzed the expression patterns of genes conferring vegetative, inflorescence and floral meristem identities. Floral repressor *FvTFL1*, conferring SAM identity, showed different expression patterns at warm and cool temperatures (**Figure 4C**). In the apices of the plants grown at warmer temperature, the expression of *FvTFL1* was lower at week 3.5 (*p = 0.0067*) and slightly lower at week 5 (*p = 0.083*), despite their later floral transition. Next, we analyzed the expression of the three woodland strawberry *LFY* homologs known to be expressed in the meristems (Zhang et al. 2023). All three *LFY* homologs were upregulated at both temperatures concordant with the floral transition but showed different temporal dynamics. Both *FvLFYa* and *FvLFYb* were strongly activated in the last time-point, with *FvLFYb* showing higher expression under cool temperature (*p = 0.011*) (**Figure 4D, E**). In contrast, the upregulation of *FvLFYa* was slightly earlier in weeks 2 (*p = 0.036*) and 3.5 (*p = 0.0085*) in the plants grown at warmer temperature, however, at week 5 the differences disappeared (*p = 0.46*). Unlike *FvLFYa* and *FvLFYb*, *FvLFYc* showed a gradual upregulation from the beginning of the treatments (**Figure 4F**). The expression of *FvLFYc* was slightly higher at week 2 (*p = 0.014*) at warmer temperature. The expression of the woodland strawberry homolog of *UNUSUAL FLOWER ORGANS* (Samach et al. 2002) *FvUFO1,* which has a conserved role in FM identity specification (Shahan et al. 2018), peaked at week 5 in both temperature treatments (**Figure 4G**). This expression pattern was similar to that of the strawberry homolog of *AP1* (**Figure 4H**) that has previously been used as a floral marker (Koskela et al. 2012). Both *FvAP1* and *FvUFO1* showed slightly higher expression at week 5 in the plants grown at cooler temperature. (*FvUFO1*, *p = 0.12*; *FvAP1*, *p = 0.068*). Moreover, the upregulation of *FvAP1* was earlier at cooler temperature (week 3.5; *p = 0.071*) corresponding to earlier floral transition.

Overall, the expression patterns of several meristem identity genes are in concordance with our phenotypic observations. However, the transcriptional dynamics of meristem identity genes (e.g. Huang et al. 2026) as well as the developmental heterochrony induced by the temperature treatments (**Figure 4A and B**) may hinder the identification of the primary regulators of temperature response.

### *FvTFL1* integrates temperature signals to fine-tune reproductive transitions in woodland strawberry

To better understand the changes in the gene expression patterns induced by the temperature treatments, we isolated the corresponding stages of SAM and FM from both temperature treatments and compared the expression of meristem identity genes. In the SAMs (1 leaf primordium initiated under treatments), *FvLFYa* was expressed higher at warmer temperature, whereas the expression of *FvLFYc* was lower. In contrast, in the FMs (sepal primordia initiated), *FvLFYb* was expressed higher at cooler temperature, whereas the expression levels of *FvLFYa*, *FvLFYc*, *FvAP1* and *FvUFO1* were similar at different temperatures (**Supplemental Figure S4A - J**). The expression of *FvTFL1* was lower at cooler temperature in the SAMs (**Figure 5A**). However, in the FMs the expression of *FvTFL1* was higher at cooler temperature (**Figure 5B**), thus correlating with the differences in the rate of floral transition and flower development. As further revealed by RNA *in situ* hybridization, *FvTFL1* mRNA was localized in the vasculature under FMs at cool temperature, whereas only weak signal was detected in the samples collected from plants grown at warmer temperature (**Figure 5C, Supplemental Figure S4 K – N**). Moreover, the expression of *FvTFL1* was very low in the secondary IMs (IM2) (**Figure 5D**). This explains why the temperature effect was mainly confined to the primary growth axis of inflorescences and the primary flowers, whereas lateral inflorescence branches and secondary flowers were unaffected by the cool temperature even when plants were exposed to treatments for eight weeks (**Figure 5E**, **Figure 2C – E, Supplemental Figure S2**). Altogether our data shows that cool temperature alters the expression pattern of *FvTFL1* leading to accelerated floral transition and delayed inflorescence and flower meristem termination, thus suggesting that *FvTFL1* integrates temperature signals to regulate timing of developmental transitions.

**Figure 5.**
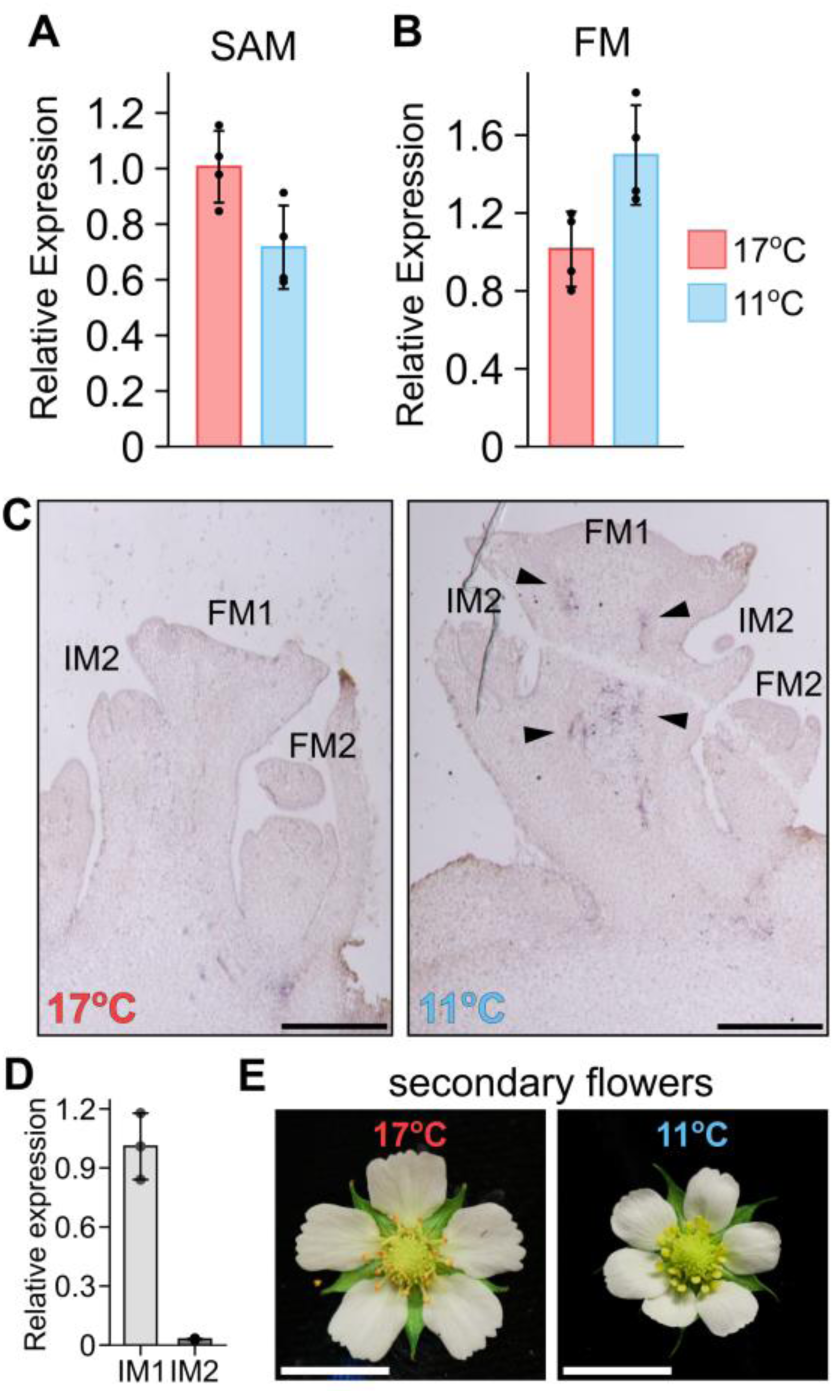
Temperature effects on reproductive transitions are associated with differences in *FvTFL1* expression. **A – B.** Relative expression of *FvTFL1* under warm (17°C) and cool (11°C) temperature treatments in the SAM with one leaf primordia initiated during the treatments (**A**) and FM with a visible sepal primordia (**B**). **C.** RNA *in situ* hybridization of *FvTFL1*. *FvTFL1* expression is visible under the primary flower meristem (FM1) at 11°C and absent at 17°C. Scale bars are equal to 250 µm. See Supplemental Figure 4 for controls. **D**. Expression levels of *FvTFL1* in the apical (primary, IM1) and lateral (secondary, IM2) inflorescence meristems. Error bars represent SDs of three biological replicates (dots). **E.** Secondary flowers of woodland strawberry showing similar numbers of flower organs when grown at different temperatures. Scale bar is equal to 10 mm.

To further test if *FvTFL1* integrates temperature signals to regulate floral transition, inflorescence branching and flower development, we used transgenic lines with silenced *FvTFL1* (Koskela et al. 2012), and *fvtfl1* mutants and wild-type plants with functional *FvTFL1* in different genetic backgrounds. WT plants with functional *FvTFL1* showed a strong response to temperatures in the timing of floral transition with a higher number of leaves before the inflorescence after the warm temperature treatment (**Figure 6A, Supplemental Figure S5A**) whereas *fvtfl1* mutants flowered with the same number of leaves after both temperature treatments, despite the differences between genotypes (**Figure 6A, Supplemental Figure S6A**). In *FvTFL1-*RNAi lines, flowering was slightly delayed at warmer temperature, but the difference was significant in only one of the lines (**Figure 6A**). Cool temperature treatment increased primary axis branching of inflorescences of *WT* plants (**Figure 6B and C, Supplemental Figure S5B**). Similarly to floral transition, silencing or loss of functional *FvTFL1* strongly reduced the effect of temperature on inflorescence branching, and in *fvtfl1* mutants, the maximum number of lateral branches was limited to two, compared with up to four branches in wild type plants grown under cool temperature (**Figure 6B & C, Supplemental Figures S5B, S6B and C).** The silencing and loss of functional *FvTFL1* also reduced the temperature-related variation in the numbers of floral organs (**Figure 6D – H, Supplemental Figure S7C and D**). The largest differences in the numbers of sepals, petals, stamens and carpels between temperature treatments were again observed in *WT* plants (**Figure 6D – H, Supplemental Figure S5C & D**). *fvtfl1* mutants produced similar number of sepals, petals and stamens under both temperatures (**Supplemental Figure S7A – E**). Interestingly, in some of *fvtfl1* mutants (H4, RdV) and secondary flowers of WT (FIN56) the number of carpels increased under cool temperatures (**Figure 6G, Supplemental Figure S3D, Supplemental Figure S6D**), suggesting that additional factors contributed to the formation of carpel primordia and termination of the flower meristem. Altogether our data indicates that *FvTFL1* integrates temperature signals to regulate the timing of developmental transitions of woodland strawberry apical meristems.

**Figure 6.**
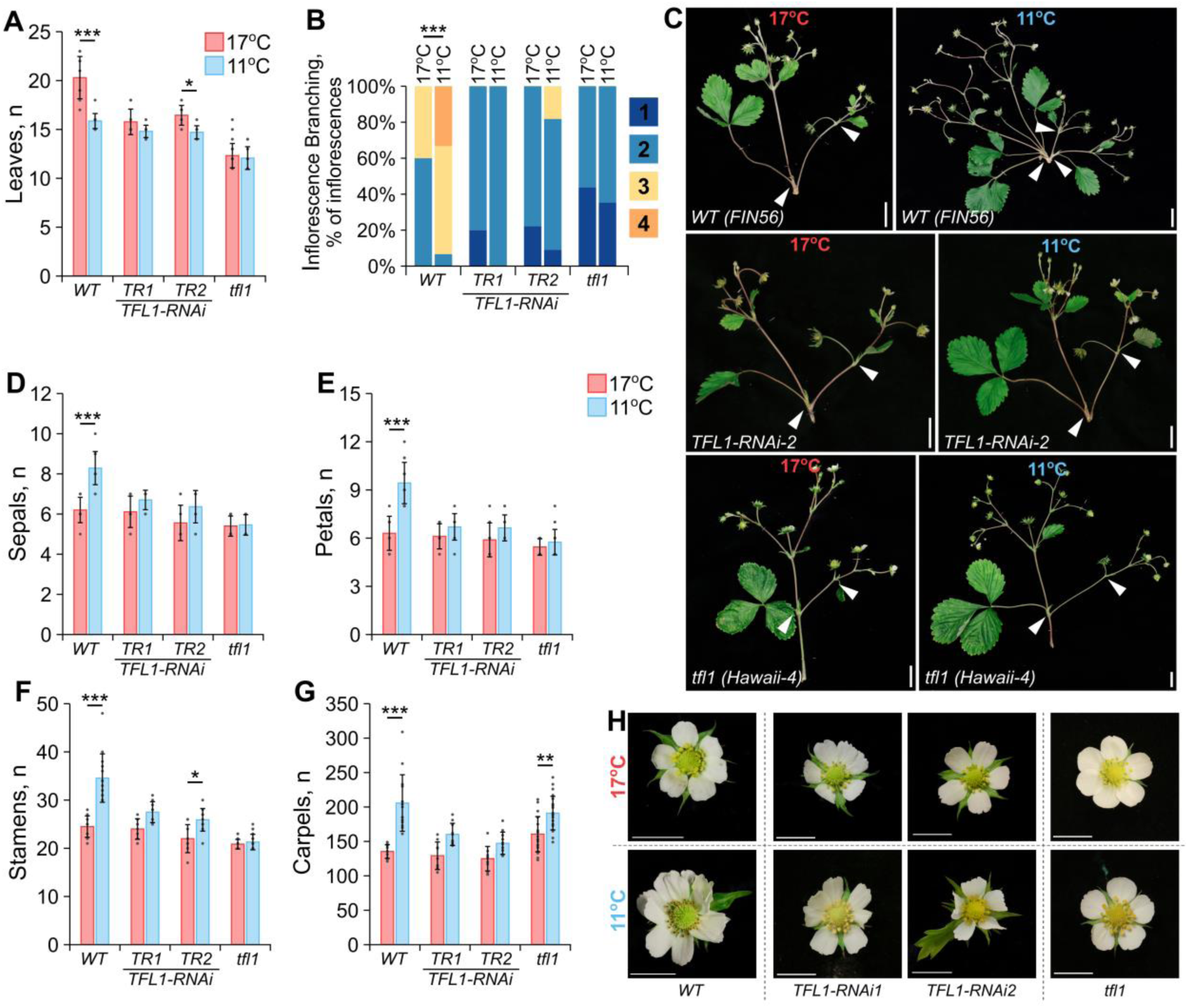
*FvTFL1* integrates temperature signals to regulate the timing of developmental transitions in woodland strawberry apical meristems. **A.** Bar plot showing the mean number of leaves until the first inflorescence (floral transition) in *WT* (FIN56), *FvTFL1*-RNAi lines (FIN56) and *fvtfl1* mutant (H4) after indicated temperature treatments of XX weeks. **B.** Bar plot showing the proportions of inflorescences with different numbers of lateral branches. **C.** Inflorescences of *WT* (FIN56) *FvTFL1*-RNAi-2, and *fvtfl1* mutant (H4) produced during indicated temperature treatments. White triangles indicate the lateral branches on the primary inflorescence axis. Scale bars are equal to 2 cm. **D – G.** Bar plots showing the mean number of sepals (**D**), petals (**E**), stamens (**F**), and carpels (**G**) on the primary flowers of woodland strawberry under indicated treatments. Error bars in **A** and **D – G** represent SDs. Dots represent biological replicates. Asterisks indicate statistically significant differences as determined by two-way ANOVA followed by Tukey’s HSD tests (* *P* < 0.05, ** *P* < 0.01, *** *P* < 0.001). **H**. Flowers of *WT* (FIN56), *FvTFL1*-RNAi- lines 1 and 2, and *fvtfl1* mutant (H4) produced during warm and cool temperature treatments. Scale bars are equal to 1 cm.

## Discussion

Using the perennial Rosaceae herb woodland strawberry as a model, we show how cool temperature dynamically modulates developmental transitions and that its effects depend on the developmental state of the meristem. The acceleration of the SAM-to-IM transition under lower temperatures contrasts with the subsequent delay in inflorescence (IM-to-FM) and flower development, indicating that the same environmental signal can have qualitatively different effects before and after floral commitment. Such stage-specific responses may be advantageous for winter annuals and perennials. Accelerating floral transition under lower temperature may allow plants to establish reproductive meristems before the onset of unfavorable winter conditions, whereas maintaining slower inflorescence and flower development could reduce the risk of exposing developing reproductive organs to damaging temperature fluctuations. However, the ecological consequences of these effects remain to be studied in detail, because, unlike photoperiod, temperature is not a stable environmental signal, and daily fluctuations can be substantial. Therefore, observations of reproductive development under fluctuating day/night and field temperatures are necessary to fully understand the effects of cool temperatures on developmental transitions. However, the described mechanism offers means to improve production of strawberries and related Rosaceae berry crops in controlled environments by reducing the duration of cropping cycles while maintaining inflorescence architecture and reproductive output.

Our results suggest that *FvTFL1* is the key regulator of the observed developmental partitioning. The temperature-dependent dynamics of *FvTFL1* expression correlate with changes in meristem identity under different temperatures, and silencing or loss-of-function of *FvTFL1* largely eliminates the temperature-dependent differences in the timing of SAM-to-IM and IM-to-FM transitions. We have previously shown that the level of *FvTFL1* expression before floral induction influences flowering time (Zhou et al. 2026). Together, these observations support a model in which both the absolute expression level and the rate of *FvTFL1* downregulation contribute to the timing of floral transition. The role of *TFL1*-like genes as repressors of reproductive transition and their gradual downregulation under low temperatures is broadly conserved across plant lineages (Jensen et al. 2001, Pillitteri et al. 2004, Wang et al. 2011, Koskela et al. 2012), however, their developmental effects can still differ between the species. For example, constitutive expression of Arabidopsis *AtTFL1* or *FvTFL1* expression in Arabidopsis delays floral transition (Ratcliffe et al. 1998; Koskela et al. 2012), whereas increased *FvTFL1* activity in strawberry completely prevents it (Koskela et al. 2012). These differences suggest that the developmental output of TFL1 activity depends on the regulatory context in which it operates. Identifying the stage-specific downstream targets of *FvTFL1* will therefore be important for understanding how it coordinates reproductive development across distinct stages and temperatures.

Woodland strawberry inflorescence combines a monopodial primary axis with sympodial lateral branches. Although the primary axis ultimately ends in a flower, it can produce a variable number of lateral branches before termination. After floral transition, *FvTFL1* continues to buffer environmental variation by regulating the timing of termination of the primary inflorescence axis. Loss of *fvtfl1* typically abolishes all but one lateral branch, a phenotype that can be partially rescued by silencing *FvFT1* (Lembinen et al. 2023). Interestingly, unlike the perennial Brassicaceae *A. alpina*, woodland strawberry does not exhibit inflorescence or flower reversion phenotypes at shorter exposures to inductive conditions (Lazaro et al. 2018), which can be attributed to the very low *FvTFL1* expression in the lateral (axillary) inflorescence meristems. The tissue specific expression pattern of *FvTFL1* not only explains why lateral inflorescence branches are unaffected by the loss of *FvTFL1* (Lembinen et al. 2023), but also why they are largely temperature insensitive. Moreover, the role of *FvTFL1* extends beyond the regulation of floral transition and inflorescence branching to regulate flower development. Unlike the timing of floral transition or termination of inflorescence meristem, the development of flowers is typically considered to be more robust and stable under environmental changes (Laitinen & Elomaa 2026). However, although the higher order flowers on strawberry inflorescences indeed appeared unaffected by temperature, the development of the primary flowers, produced by apical meristems, was severely affected at cool temperature. Overall, differential regulation of *FvTFL1* enables a balance between environmental plasticity and developmental robustness, allowing variation in the number of lateral branches and number of organs on primary flowers while maintaining stable branching patterns and robust development of flowers in higher-order axes.

Importantly, *FvTFL1* is unlikely to function as a temperature sensor itself. Rather, its stage-specific expression dynamics may represent an integration point through which upstream temperature-responsive pathways alter meristem development. However, despite the central role of *TFL1* genes in the regulation of flowering in strawberry and other Rosaceae species (Koskela et al. 2012, Iwata et al. 2012, Freiman et al. 2012, Flachowsky et al. 2012, Koskela et al. 2016, Bai et al. 2017, Fan et al. 2022), their upstream regulation remain largely unresolved. Recent evidence from woodland strawberry indicates that *FvFT1* maintains *FvTFL1* expression in the shoot apical meristem under long-day conditions, thereby preventing floral transition during summer (Zhou et al. 2026). However, the expression of *FvFT1* is suppressed under short days indicating that its contribution to *TFL1* regulation in autumn is limited. Recent results from Arabidopsis highlight the importance of a negative feedback loop between *LFY* and *TFL1* in regulating inflorescence development (Huang et al. 2026). In IMs, LFY promotes the expression of *TFL1*, thus helping to maintain their indeterminacy. Unlike in Arabidopsis, however, *FvTFL1* expression in strawberry declines already during the transition from SAM to IM, an expression pattern opposite to that of each of the three *LFY* homologs in woodland strawberry. Recent functional studies of the woodland strawberry *LFY* homologs have demonstrated that *FvLFYa* has a conserved role in inflorescence and flower development (Zhang et al. 2023). However, although *FvLFYb* can complement Arabidopsis *lfy* mutant, its specific role in woodland strawberry remains elusive. Interestingly, the expression of *FvLFYb* was higher in FMs at cool compared with warm temperature, suggesting that it may promote *FvTFL1* expression. In contrast, loss of *FvLFYc* had no apparent effect on inflorescence or flower development (Zhang et al. 2023). Importantly, both *FvLFYa* and *FvLFYc* have been characterized only in the *fvtfl1* mutant background, leaving the potential regulatory interactions between strawberry *FvTFL1* and *FvLFY* orthologs unresolved.

Altogether, our findings establish *FvTFL1* as a regulator linking temperature responses to reproductive transitions. Rather than acting as an on–off regulator of flowering, *FvTFL1* integrates developmental stage and meristem identity to modulate the timing of floral transition, inflorescence termination, and, to a lesser extent, flower development, thereby balancing developmental plasticity with robustness. The distinct temperature responses of successive meristem stages further indicate that reproductive development is not uniformly regulated by temperature but instead involves stage-specific regulatory mechanisms.

## Materials and Methods

### Plant material and growth conditions

Woodland strawberry (*Fragaria vesca* L.) accession FIN56 was originally obtained from the National Clonal Germplasm Repository, Corvallis (PI551792) and maintained through self-pollinations. To investigate the role of *FvTFL1* in regulating temperature dependent developmental transitions we used *FvTFL1-*RNAi lines in FIN56 background (Koskela et al. 2012) and five *F. vesca semperflorens* accessions: Hawaii-4 (H4), Yellow Wonder (YW), Reine des Valles (RdV), Baranovka and Piikkiö, all carrying a 2-bp deletion in the first exon (Lembinen et al. 2023). Experiments were performed on either runner-propagated or seed-germinated plants.

For the assessment of temperature effects on plant development, timing of meristem identity transitions, and gene expression, we used runner-propagated FIN56 plants. Runner cuttings were rooted on peat tablets (Jiffy Products International BV, Netherlands) and pre-grown in the greenhouse at 18°C and 18h/6h day/night photoperiod. Plants were illuminated with 150 µmol m^-2^ s^-1^ light intensity using high-pressure sodium lamps (Airam 400W, Finland). After the initial pre-growth of four weeks, plants were transferred into 10x10 cm pots filled with peat substrate (Kekkilä, Finland). Plants were then moved into controlled climate growth rooms and grown there for another week at 18°C and 18h/6h day/night photoperiod before the start of treatments. During treatments, the conditions in the growth rooms were set to either 11°C (cool) or 17°C (warm) and 12h/12h day/night photoperiod. In the growth rooms plants were illuminated with LED lamps (Valoya AP67, Finland), with ∼150 µmol m⁻² s⁻¹ light intensity. After the treatments plants were moved back to greenhouse for phenotypic observations (20°C and 18h/6h day/night). Plants were irrigated with tap water supplemented with fertilizers (N-P-K 13-7-20; GreenCare) 0.3 g/l and Calcinit 0.1 g/l (YARA, Norway).

To investigate the role of *FvTFL1* in the temperature dependent regulation of developmental transitions we first performed an experiment using runner-propagated *FIN56* (*FvTFL1*), *NOR8* (*FvTFL1*) and *Hawaii-4* (*fvtfl1*). Plants were pre-grown in the greenhouse as previously described. *Hawaii-4* (*fvtfl1)* runner plants developed multiple inflorescences prior to the treatments. To ensure that the observations of the temperature effect are done on the inflorescences and flowers developed under the treatment conditions, we removed all the inflorescences developed during pre-growth and the first five inflorescences developed during the treatments. *FIN56* and *NOR8* plants were grown under the treatments for 7 weeks. *Hawaii-4* plants were kept in the treatments for three months.

To include the additional *fvtfl1* mutants (Yellow Wonder, Reine des Valles, Baranovka, Piikkiö), we conducted additional experiments using seeds as starting material. Seeds of *FIN56*, *FvTFL1*-RNAi transgenic lines, and five *fvtfl1* mutants were germinated on wet filter paper at 22°C and 18h/6h photoperiod. Germinated seeds were transferred into peat tablets (Jiffy Products International BV, Netherlands) at the cotyledon stage and pre-grown in growth room at 18°C and 18h/6h day/night photoperiod until the appearance of the second juvenile leaf. After that the seedlings were separated into treatments. During the treatments, when all plants, within a designated treatment, developed at least 6 true leaves they were transferred into 10x10 cm pots filled with peat substrate (Kekkilä, Finland). Plants were grown under the treatment temperatures for three months until FIN56 plants completed their flower meristem development which was assessed under stereomicroscope. By that time, all *fvtfl1* plants and some of *FvTFL1*-RNAi plants already had visible flower buds. Plants were then transferred to greenhouse for phenotypic observations.

### Phenotyping and statistical analysis

Flowering time was recorded as the number of days after the end of the treatment or the number of leaves until the first inflorescence. In runner-propagated plants, the youngest fully developed leaf was marked in the beginning of the treatment, whereas in seed-derived material, counting started from the first leaf after cotyledons. The inflorescences produced on the main rosette were analyzed. The inflorescence branching was evaluated by counting the number of branches produced along the primary growth axis. The total number of flowers per inflorescence was counted. The number of sepals and petals was counted from the primary or secondary flowers at the anthesis. Numbers of stamens and carpels were counted from the photographed flowers using Fiji ImageJ (Schindelin et al. 2012). Photos were taken using Canon 800D camera with macro lens EF 100mm 1:2.8 (Canon, Japan) from flowers at 5 – 10 days after anthesis to ensure high level of contrast between carpels. The ripen berries formed by primary flowers were collected and their fresh weight measured. The berries were then photographed, and their width was measured in Fiji ImageJ. For figure preparation, photo cropping, white-balance, and background removal was done using Photoshop CC 2019 (Adobe, US).

All data was analyzed using one- or two-way ANOVA or Student’s *t*-test as indicated in figure legends. Significant differences between treatments and/or lines at *P* < 0.05 were determined using Tukey’s HSD multiple comparisons tests. All analyses were performed in R v4.3.0 (R Core Team 2023).

### RNA extraction and RT-qPCR

For time course expression analysis, shoot apex samples were collected under the treatments at time points indicated in the figures or figure legends and stored at -80°C before carrying out RNA extraction. Each biological replicate consisted of three shoot apices. For the analysis of the stage specific expression, we collected SAMs after development of one leaf primordia under the treatments and FMs at the sepal primordia stage. The meristems were dissected under stereomicroscope. Each biological replicate consisted of 5 – 7 meristems.

Total RNA was extracted as in Mouhu *et al*. (2009). Protoscript II reverse transcriptase kit (NEB, US) was used for cDNA synthesis. qRT-PCR was performed using a LightCycler 480 SYBR Green I Master kit in a Roche LightCycler 480 (Roche Diagnostics, Switzerland), as described in Mouhu et al. (2009). Each gene was tested by three technical replicates. The expression was calibrated using *FvMSI1 (FvH4_7g08380*) and *FvUBQ_10_* (*FvH4_3g02641*) as reference genes. Primer sequences used in this study are shown in **Supplemental Table S1.**

### RNA in situ hybridization

For RNA in situ hybridization, we used full length probes amplified from cDNA from SAM using reverse transcription-PCR. The probes were amplified with primers containing a T7 overhang at the 5’ end of the forward (sense probe) and the reverse (anti-sense probe) primer. The primer sequences are listed in **Supplemental Table S1**. The gel-purified PCR products were used as templates for in vitro transcription with T7 RNA polymerase and labeled with DIG RNA Labeling Kit (Roche, Switzerland) according to the instructions. The labeled probes were hydrolyzed to approximately 150bp as in Karlgren et al. (2009).

Apex samples containing the primary floral meristem were dissected from the wild-type (FIN56) plants grown under 11 and 17°C. The samples were fixed in FAA solution (3.7% [v/v] formaldehyde, 5% [v/v] acetic acid, 50% [v/v] ethanol), dehydrated using ethanol series, cleared in TissueClear (Sakura Finetek, Japan), and embedded in paraffin (Sigma-Aldrich, US). Sectioning was done using Rotary Microtome Microm HM 355S (Thermo Fisher Scientific, US). Hybridizations were done as described in Koskela et al. (2012). Sections were photographed using a Leica M205 FCA microscope equipped with a Leica DMC6200 camera (Leica Mikrosysteme Vertrieb GmbH, Germany).

### SEM imaging

Meristem samples were dissected from wild type plants (FIN56), at weekly intervals. The samples were fixed, dehydrated, critical point dried, and coated with platinum as previously described (Lembinen et al. 2023). The imaging was performed using a Quanta 250 SEM (FEI Corporation, US) at the Electron Microscopy Unit, Institute of Biotechnology, University of Helsinki.

### Accession numbers

Sequence data from this article can be found in the *Fragaria vesca* (ver4.0) genome under accession numbers: *FvTFL1* FvH4_6g18480, *FvLFYa* FvH4_5g09660, *FvLFYb* FvH4_3g04170, *FvLFYc* FvH4_3g03530, *FvUFO1* FvH4_3g12960, *FvAP1* FvH4_4g29600, *FvMSI1* FvH4_7g08380, *FvUBQ10* FvH4_3g02641.

## Acknowledgements

We thank present and former members of the Hytönen laboratory for discussions and input on the manuscript; Eija Takala for laboratory assistance and Katriina Palm for plant maintenance. The authors acknowledge the support provided by the Electron Microscopy Unit of the University of Helsinki. This work was supported by the Research Council of Finland (Grants No. 317306 and 368864 to TH) and Jane and Aatos Erkko Foundation (grant 4709012 to PE and TH).

## Author contributions

SL and TH designed the research. SL, QZ and GF performed the experimental work. FW performed RNA *in situ* hybridization. TZ contributed to experimental design and data analysis. TH and PE supervised the work. SL and TH wrote the manuscript with contributions from all authors.

