## Supplemental Data for "TFL1 homolog integrates temperature signals to control timing of developmental transitions in woodland strawberry"

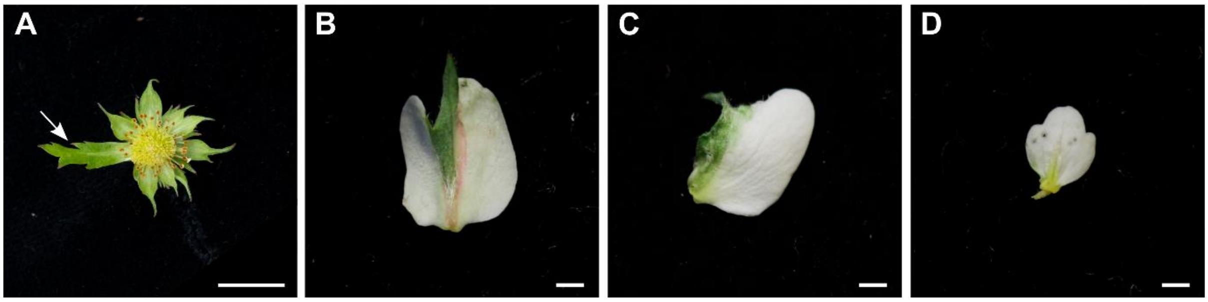


**Supplemental Figure S1. Chimeric floral organs on the primary flowers of woodland strawberry. A.** Bract-like sepal. Scale bar is equal to 1 cm **B – C**. Sepaloid petals. **C**. Petaloid stamen. Scale bars are equal to 1 mm.

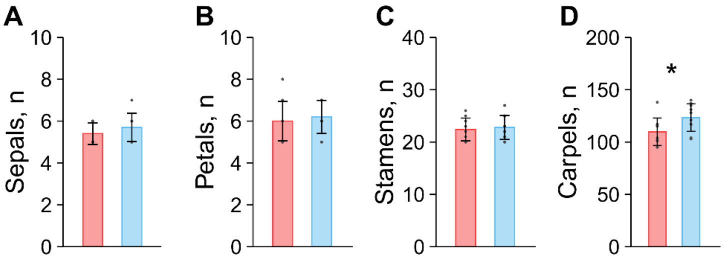


**Supplemental Figure S2. Effect of temperature on the development of the higher order (secondary) flowers of woodland strawberry.** Bar plots showing the mean numbers of sepals (**A**), petals (**B**), stamens (**C**), and carpels (**D**) on the flowers produced at 17°C (red) and 11°C (blue). Error bars represent SDs of 10 replicates (dots). Asterisk indicates statistically significant differences as determined by Student's *t*-test (* $P < 0.05$).

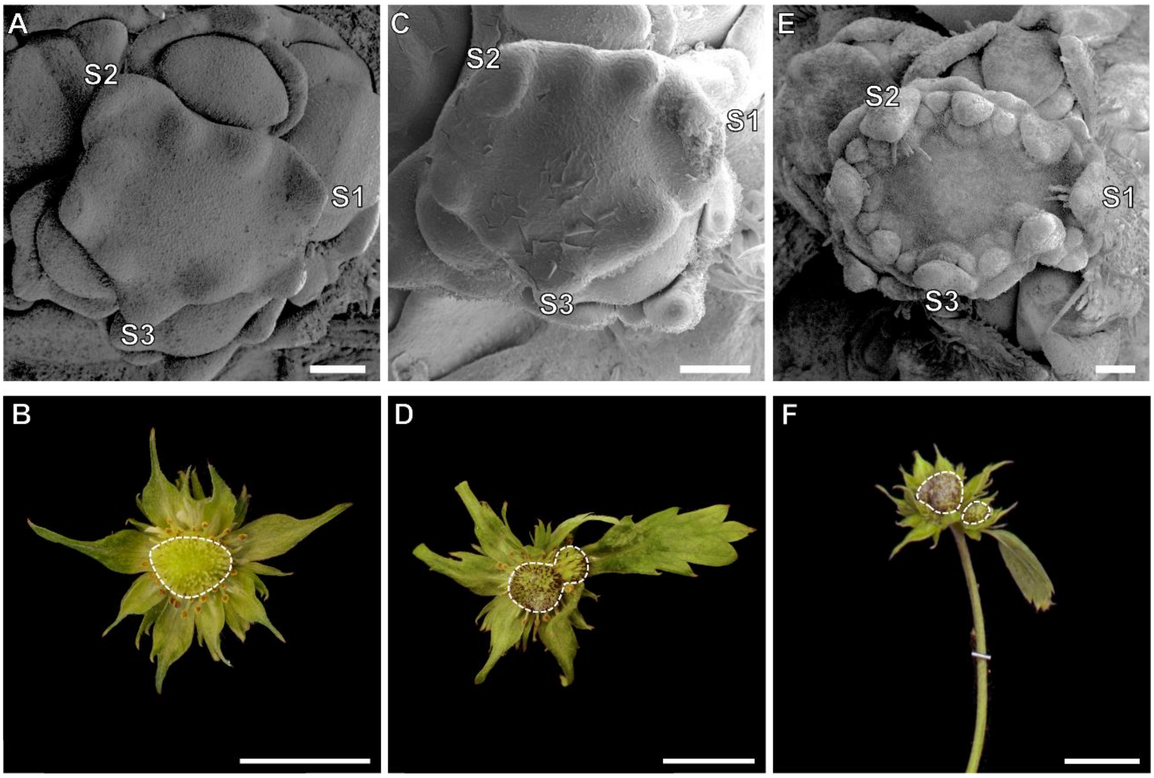


**Supplemental Figure S3. Receptacle fasciation phenotypes**. **A.** SEM image showing the how the first three sepals (S1 – S3) grow farther from the meristem. **B.** Mild fasciation of the receptacle stretching toward the first three sepals. **C.** SEM image showing larger gap between S1 and meristem which may lead to formation of the larger fasciation. **D.** Intermediate fasciation of the receptacle, with the longer receptacle. **E.** SEM image showing further separation of S1 from the meristem which may lead to formation of a boundary and separation of the receptacle. **F.** The severe fasciation with two receptacles separated by sepals. Scale bars in A, C, D are equal to 100 µm. Scale bars in B, D, F are equal to 1 cm.

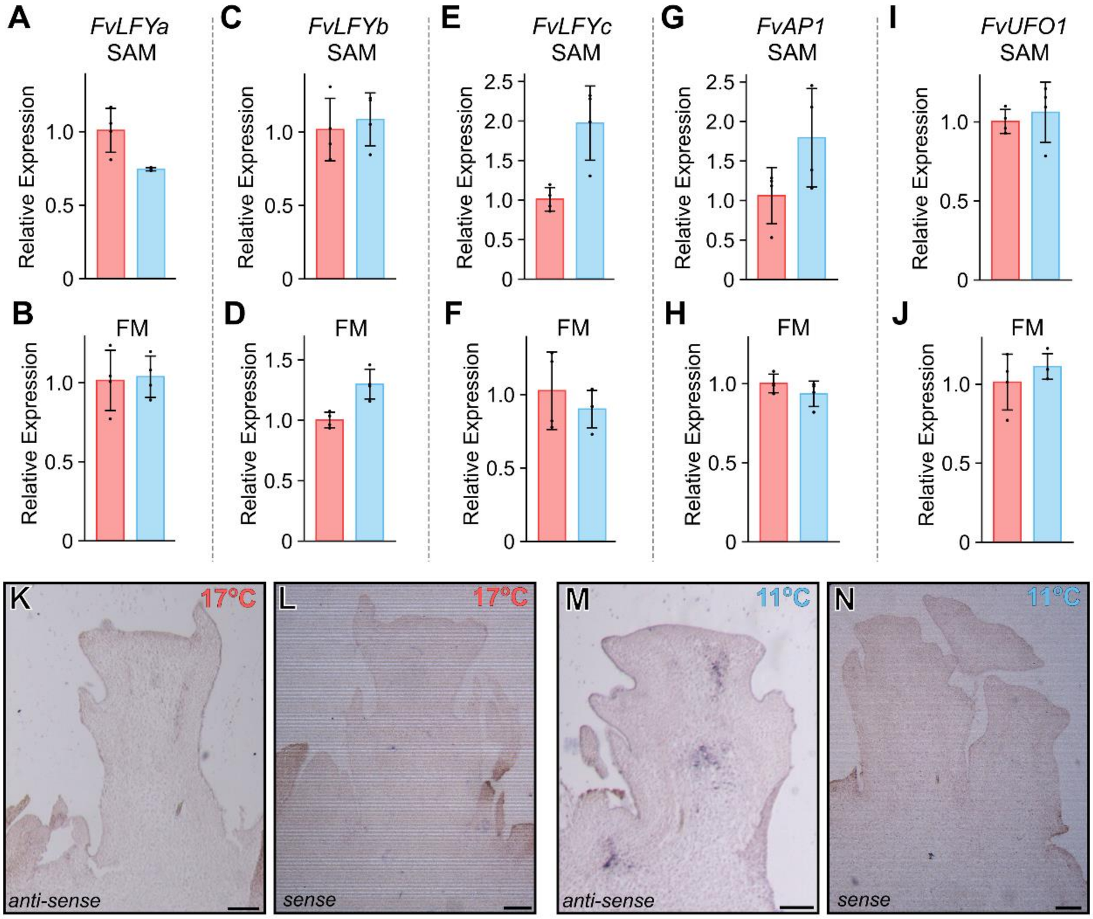


**Supplemental Figure S4. Expression patterns of meristem identity genes in the shoot apical (SAM) and flower (FM) meristems. A – J.** Bar plots showing the mean expression levels of *FvLFYa* (**A, B**), *FvLFYb* (**C, D**), *FvLFYc* (**E, F**), *FvAP1* (**G, H**), *FvUFO1* (**I, J**) at warmer (red) and cooler (blue) temperatures. Error bars represent SDs of three to four biological replicates (dots). **K – N.** RNA *in situ* hybrization of *FvTFL1* anti-sense and sense probes in samples from warm (**K, L**) and cool temperature (**M, N**).

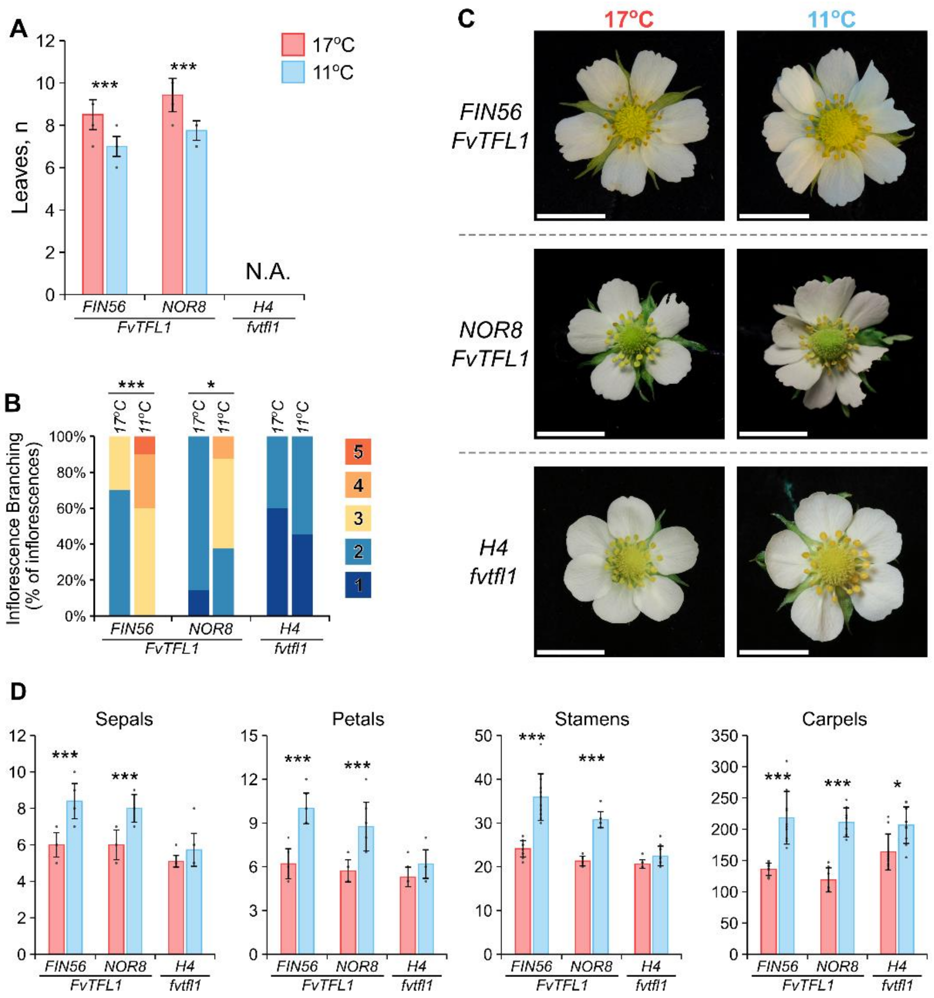


**Supplemental Figure S5. Temperature effect on the developmental transitions in runner propagated wild-type (FIN56 and NOR8) and *fvtfl1* mutant (H4) plants. A.** Bar plot showing the mean number of leaves until the first inflorescence. **B.** Bar plot showing the proportions of inflorescences with different numbers of lateral branches. **C.** Photos showing the flowers WT (*FIN56* and *NOR8*), and fvtfl1 (*H4*) plants developed during the temperature treatments. Scale bars are equal to 1 cm. **D**. Bar plots showing the mean numbers (y-axis) of sepals, petals, stamens and carpels. Error bars in A and D represent SDs of 8 – 10 replicates (dots). Asterisks indicate statistically significant differences as determined by two-way ANOVA followed by Tukey's HSD tests (* $P < 0.05$, ** $P < 0.01$, *** $P < 0.001$).

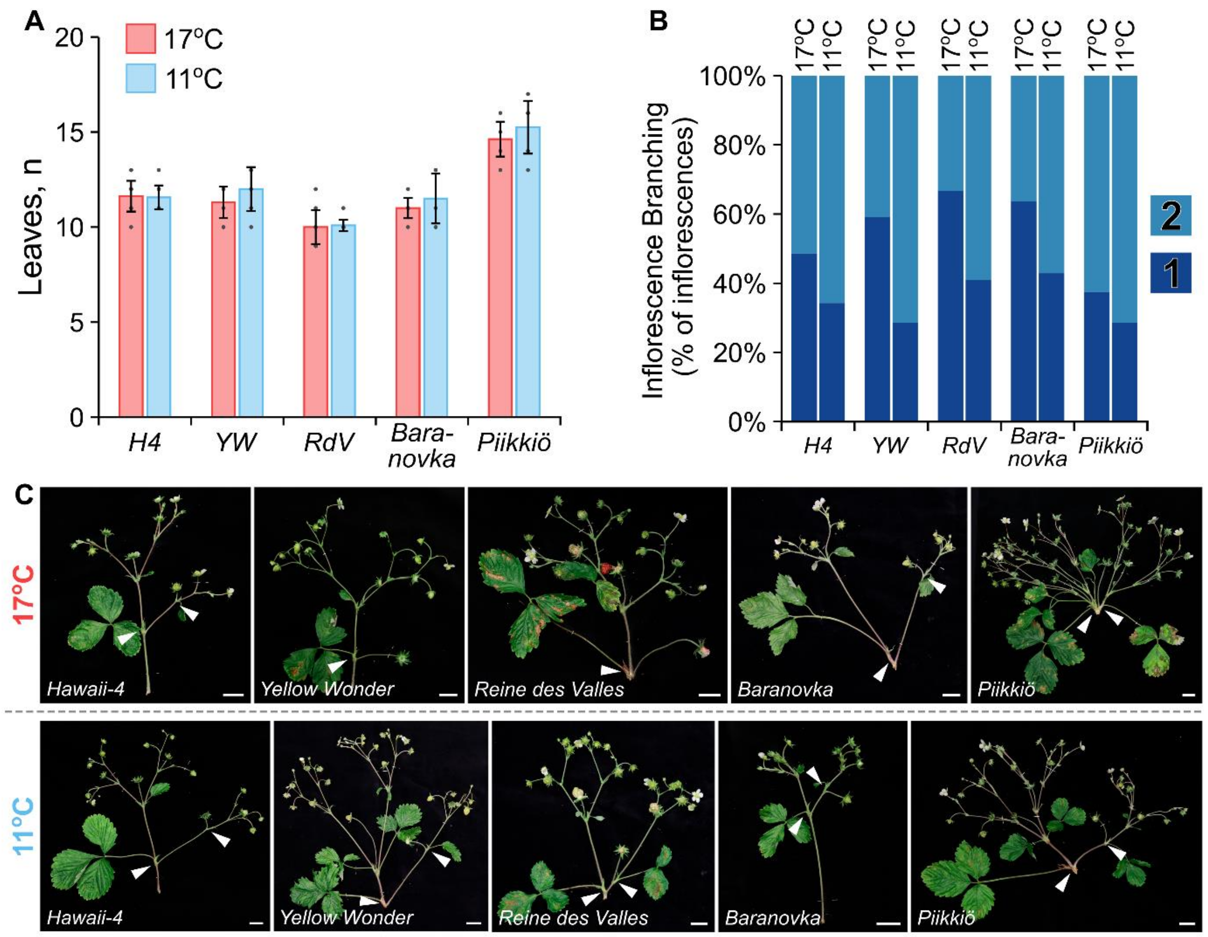


**Supplemental Figure S6. Temperature effect on floral transition and inflorescence development in *fvtfl1* mutants in different backgrounds. A.** Bar plot showing the mean number of leaves until the first inflorescence. Error bars represent SDs. Dots represent individual plants (n = 8 – 16). **B.** Bar plot showing the proportions of inflorescences with different numbers of lateral branches. **C**. Inflorescence photos of *fvtfl1* mutants in different backgrounds produced during the temperature treatments showing the lateral branches (white triangles) on the primary growth axis. Scale bars are equal to 2 cm.

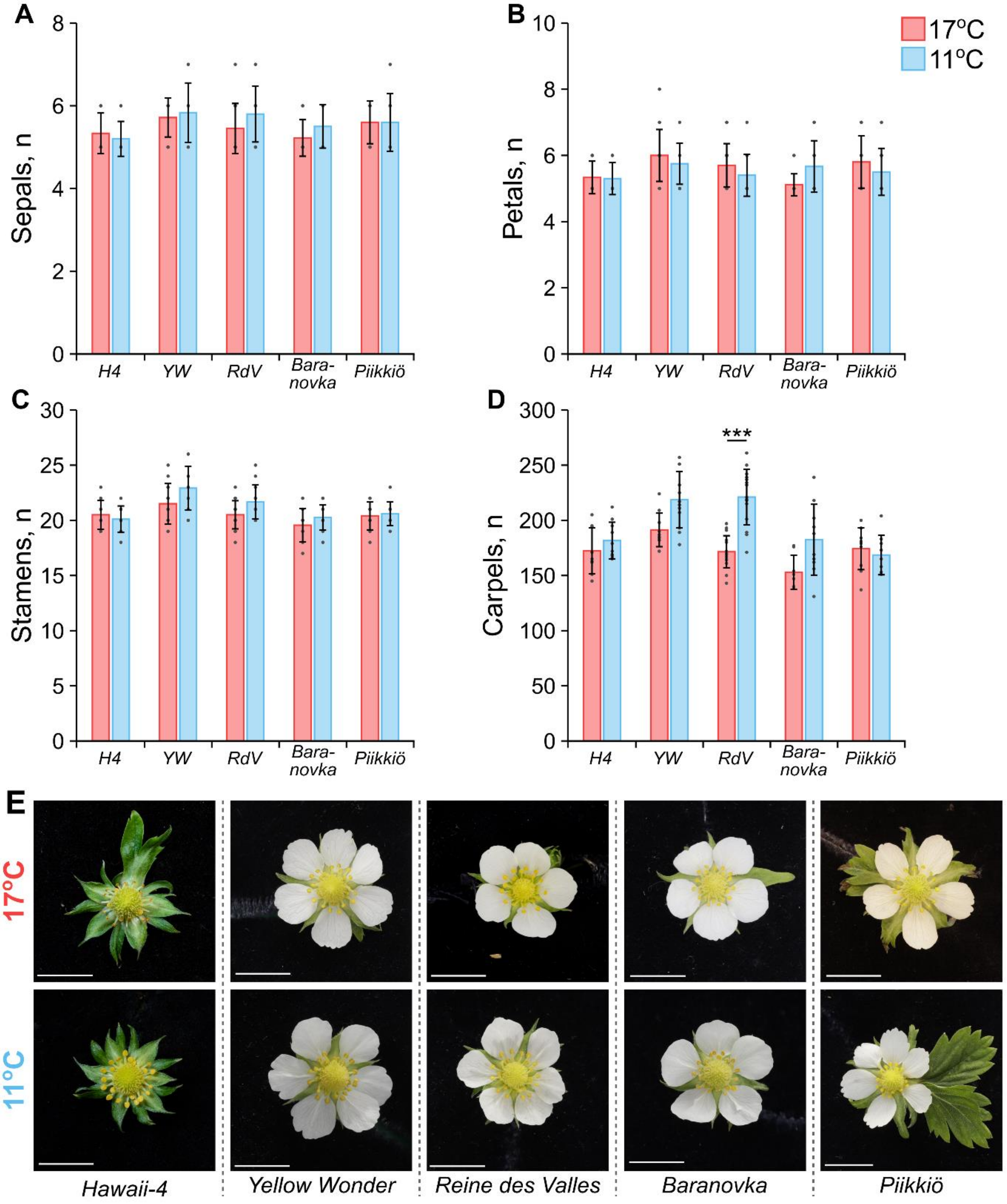


**Supplemental Figure S7. Temperature effect on flower development in *fvtfl1* mutants in different backgrounds. A – D.** Bar plots showing the mean numbers of sepals (**A**), petals (**B**), stamens (**C**), and carpels (**D**) on the flowers produced during the temperature treatments. Error bars represent SDs (n = 8 – 15, dots). Asterisk indicates statistically significant differences as determined by two-way ANOVA followed by Tukey's HSD tests (*** $P < 0.001$). **E**. Flower photos of five *fvtfl1* mutants from two temperature treatments. Scale bars are equal to 1 cm.

**Supplemental Table S1.** Primers used in experiments.

| Primer name | Sequence (5' - 3') |
|---|---|
| | **RT-qPCR** |
| FvTFL1-F | AACGGCAGCAACAGGAAC |
| FvTFL1-R | CTGGCACCACAGATGCTACA |
| FvUFO1-F | AGGCTCCCATGAGAAGGTTT |
| FvUFO1-R | CCGTTTCCATTTTCAAGTGTTG |
| FvMSI1-F | ACACCATCAGTCTCCTGCCAAG |
| FvMSI1-R | TCCCCACACCTTTGATTGCCA |
| FvUBQ10-F | CAGACCAGCAGATTATCTT |
| FvUBQ10-R | TTCTGGATATTGTAGTCTGCTAGGG |
| FvAP1-F | AGCTCAGGAGGTTCATGACTG |
| FvAP1-R | TAAGGTCGAGCTGGTTCCTC |
| FvLFYa-F | GATGACAGAATCAATGGAGGAG |
| FvLFYa-R | CTGGTTTGTGACCTTGGTGG |
| FvLFYb-F | AACACTGCCAAGGAGCGTGGT |
| FvLFYb-R | CGCGTAGCAGTGAACGTAATG |
| FvLFYc-F | GAGACGACGACTTGAGGAGGA |
| FvLFYc-R | CCGCTTCTTCGCCGCCCTAA |
| | **RNA *in situ*** |
| FvTFL1-F | ATGGCAAGAATGTCGGAACC |
| FvTFL1-T7-R | cataatacgactcactatagggCTAGCGTCTTCTTGCTGCC |
| FvTFL1-T7-F | cataatacgactcactatagggATGGCAAGAATGTCGGAACC |
| FvTFL1-R | CTAGCGTCTTCTTGCTGCC |
